# Neural plate targeting by in utero nanoinjection (NEPTUNE) reveals a role for *Sptbn2* in neurulation and abdominal wall closure

**DOI:** 10.1101/2020.05.28.115972

**Authors:** Katrin Mangold, Jan Mašek, Jingyan He, Urban Lendahl, Elaine Fuchs, Emma R Andersson

## Abstract

Gene variants associated with disease are efficiently identified with whole genome sequencing or GWAS, but validation *in vivo* lags behind. We developed NEPTUNE (<u>ne</u>ural <u>p</u>late targeting by in <u>u</u>tero <u>n</u>anoinj<u>e</u>ction), to rapidly and flexibly introduce gene expression-modifying viruses to the embryonic murine neural plate prior to neurulation, to target the future adult nervous system. Stable integration in >95% of cells in the brain enabled long-term gain- or loss-of-function, and conditional expression was achieved using mini-promotors for cell types of interest. Using NEPTUNE, we silenced *Sptbn2*, a gene associated with Spinocerebellar ataxia type 5 (SCA5) in humans. Silencing of *Sptbn2* induced severe neural tube defects and embryo resorption, suggesting that *SPTBN2* in-frame and missense deletions in SCA5 reflect hypomorphic or neomorphic functions, not loss of function. In conclusion, NEPTUNE offers a novel, rapid and cost-effective technique to test gene function in brain development, and can reveal loss of function phenotypes incompatible with life.

## Introduction

Recent progress in genomics, metagenomics and transcriptomics has facilitated rapid identification of gene variants associated with diseases in humans (*1, 2*). *In vivo* validation in mammalian systems has, however, lagged behind since generation of novel mouse models is limited by the long reproduction time of mice. Previous work using ultrasound-guided *in utero* transduction with fluorescently traceable lentiviruses, carrying RNAi or Cre recombinase, into mouse embryos demonstrated noninvasive and highly efficient transduction of surface epithelium (*3, 4*) allowing rapid deciphering of gene networks and mechanisms in skin (*5, 6*). In contrast, genetic manipulation of developing mouse brain has largely relied on *in utero* electroporation (*7, 8*) or viral infection (*9*) post-neurulation, which targets a smaller subset of cells in the brain and cannot achieve long-term and wide-spread gene manipulation, nor systemic effects. Pre-neurulation injection of virus or fluorescent beads into amniotic fluid is tolerated by mouse embryos and can be used for targeting of the future brain (*10, 11*). We therefore developed NEPTUNE (<u>Ne</u>ural <u>P</u>late targeting by in <u>u</u>tero <u>n</u>ano-inj<u>e</u>ction), to rapidly and flexibly introduce gene expression-modifying viruses to the neural plate prior to neurulation and achieve widespread or conditional stable transduction, in order to test gene function rapidly and efficiently during nervous system development. Commercially available validated shRNA viral vectors can be used to generate virus (three days for production) and after in utero transduction at E7.5, collection of tissue depends on stages of interest. The time from hypothesis to results is thus dramatically shortened compared to generating knockout mice.

Mutations in *SPTBN2* (encoding the membrane scaffold protein β-III SPECTRIN) are associated with multiple human nervous system disorders. Heterozygous in-frame deletions or missense mutations have been identified in progressive autosomal-dominant adult-onset Spinocerebellar Ataxia Type 5 (SCA5) (*12*–*16*) while homozygous mutations in *SPTBN2* are associated with SCAR14 (spinocerebellar ataxia, autosomal recessive 14, also known as Spectrin-associated Autosomal Recessive Cerebellar Ataxia type 1 (SPARCA1)) (*17*–*19*). However, de novo point mutation of *SPTBN2* has also been described in ataxic cerebral palsy (*20*). In mice, homozygous *Sptbn2* targeting recapitulates adult-onset progressive ataxia, but persistent expression of shorter SPTBN2 isoforms (*21*) or expression of alternative splice-variants (*22*) suggest that this phenotype may reflect hypomorphism or neomorphism rather than complete loss of function. Development of a rapid technology to manipulate gene expression in the mouse nervous system would allow testing of the function of *Sptbn2* during embryonic development, while bypassing the risk of isoform expression or genetic compensation (*23*).

In this study, we developed a new method to manipulate gene expression in the developing murine nervous system, which we dubbed <u>ne</u>ural <u>p</u>late targeting by *in <u>u</u>tero* <u>n</u>anoinj<u>e</u>ction (NEPTUNE). Taking advantage of the accessibility of the pre-neurulation neural plate, we demonstrated that *in utero* injection at this stage results in wide-spread transduction of the future brain and spinal cord, enabling long-term gain- and loss-of-function experiments. In order to achieve conditional expression while avoiding the use of dedicated Cre mouse strains, we used previously identified mini-Promotors to drive cell-type specific expression (*24*) in neuronal progenitors, astrocytes, and oligodendrocytes. Finally, we tested the function of *Sptbn2* in nervous system development to assess whether NEPTUNE can achieve systemic effects. Our results show that NEPTUNE is a powerful new method for manipulation of gene expression in the developing nervous system achieving near 100% transduction of the brain, which can be coupled with promotors to achieve cell-type specific expression, and revealing new functions for human disease-associated genes.

## Methods

### Animals

CD1 wild type mice were obtained from Charles River Laboratories. Animals were housed according to European regulations, with a standard day and night cycle with food and water ad libitum. Females were checked for estrus and naturally mated overnight. 8 week old females were plugged overnight, and gestation was defined as embryonic day (E) 0.5 at noon of the same day of vaginal plug. Ethical approval for all experiments described here was granted by the Swedish Board of Agriculture (Jordbruksverket) with permit numbers N59/14 and 8188-2017.

### Ultrasound check of pregnancy and anesthesia

Pregnancy in plug positive females was confirmed via ultrasound (US) the day before injection. The pregnant female was placed in an induction box and anesthetized with an initial dose of 3-4% Isoflurane (000890, Baxter Medical AB). Once anesthetized, the female was moved to a heated surgical table, pre-warmed to 37°C. In order to maintain anesthesia, the snout was placed into an attached nose cone and the Isoflurane dose was lowered to 1.5-2%. All four paws were fixed with surgical tape and the hair around the lower abdomen was removed using commercial Veet Hair Removal cream. Ultrasound gel was applied to the clean dry skin, and the presence of embryos was assessed using US. Afterwards the belly was wiped clean with water and the mouse was placed back into its cage. Awakening was monitored and an additional check was performed 15 minutes later.

### Surgery and ultrasound-guided nanoinjections

For anesthesia and placement for surgery the same routines as for US-check of pregnancy were used. Once asleep, the eyes were covered with commercial eye gel (APL 3044285,) to prevent drying of the eyes, and 0.1 mg/kg Buprenofine (521634, Indivior Europe Limited) was injected sub-cutaneously. With a pair of surgical scissors, a 1-2 cm vertical midline incision was made to open the lower abdomen. Using two pairs of surgical forceps, carefully both uterine horns were exposed, and the total number of embryos recorded. For the injections, the top embryos of the left or right uterine horn were left exposed, while the remaining embryos were carefully pushed back into the abdominal cavity with a sterile cotton tip. The embryos as well as the surrounding skin were re-hydrated with sterile PBS. Next, 3-4 embryos were pulled through the elastic bottom of a modified petri dish, which was secured on the surgical table using commercial play dough. The dish was then filled with sterile PBS until the embryos were immersed. In order to prevent leakage, the elastic bottom can be pushed down with a cotton tip, so that it adheres to the wet, surrounding skin. The embryos in the petri dish were further immobilized with an additional piece of play dough, to avoid any unwanted movement during injection. The US-probe was lowered into the PBS and the stage moved to the first embryo. The glass needle (in-house pulled capillary, grinded tip), loaded with virus and attached to a nanoinjector (W369-0131; Harvard Apparatus), was also lowered into the PBS and via US aligned to the embryo.

Following the uterine horn, the next 3-4 embryos were pulled through the elastic while the already injected ones were carefully put back. After surgery and injection, the muscular layer of the abdomen was sutured using 6-0 prolene (J384H, Angthos) and the skin was clipped with EZ-clips (59027, Angthos). The female was put back into a pre-warmed cage and recovery was carefully monitored, with an additional check 15-30 minutes after surgery. All work was carried out in a laminar air flow hood. In between mice, all surgical instruments (forceps, scissors etc.) were sterilized in a 200°C glass bead sterilizer. Disposable products (cotton tips, tissues) were replaced with fresh ones.

### Tissue collection and fixation

Females were sacrificed in a CO^2^ chamber. Whole uterine horns were exposed, and embryos dissected under a microscope in sterile, room temperature PBS. Tissues were fixed overnight at 4°C in 4% formalin diluted in sterile PBS, followed by washing in PBS and dehydration in 30% sucrose buffer. Samples were embedded in OCT freezing medium and stored at −80°C. 12µm cryosections were prepared at −20°C at a cryotome. Sections were kept at −20°C or −80°C for short- and long-term storage, respectively.

### Immunohistochemistry

Sections were thawed at room temperature and the edges of the glass slides were outlined with a PAP Pen to create a hydrophobic barrier. Slides were re-hydrated in PBS, followed by blocking for 1h at room temperature (5% Donkey Serum (D9663, Sigma) in 0.3% PBS-Tween (Tween 20; P9416 from Sigma and PBS tablets; 003002 from Invitrogen)). Primary Antibody was incubated overnight at 4°C in the dark (prim Ab in 0.3% PBST). The next day, slides were rinsed in PBS and incubated in secondary Antibody for 1 hour at room temperature (secondary Antibody in blocking solution). After an additional washing, slides were mounted using ProGlo mounting medium (P36961, Thermo Fisher) and stored at 4°C. Sections were imaged using confocal LSM880 microscope. Primary Antibodies that were used: goat anti-Sox2 (sc-1723, Santa Cruz; 1:200); chicken anti-GFP (ab13970, Abcam; 1:1000); mouse anti-NeuN (MAB 377, Merck 1:200); rabbit anti-Gfap (Z0334, Agilent; 1:1000); goat-anti Olig1 (MAB2417, R&D systems 1:100); guinea pig anti-Dcx (AB2253, Merck 1:250).

### Virus production and titration

Low passage (<p10) Lenti-X™ 293T cells (632180, Clontech) were cultured in DMEM High Glucose (41965039, Invitrogen; 10131-027 Invitrogen), supplemented with 10%FBS (10270106, Invitrogen), 1%Pen/Strep and 1% Geneticin (15140122, Invitrogen). 24 hours prior to transfection, cells were seeded onto two 500 cm^2^ plates (2 plates per virus), that were pre-coated with 0.01mg/mL in PBS Poly-L-Lysine (P5899, Sigma). Cells were transfected using the calcium phosphate transfection method: 275µg of vector plasmid, 275µg of psPAX2 (packaging plasmid) and 180µg of pMD2.G (VSV-G plasmid) were mixed in a 50mL conical tube. (pMD2.G and psPAX2 were a gift from Didier Trono (Addgene plasmid #12259 and #12260). 2.28mL 2M CaCl_2_ (BP9742, Fisher Scientific) and MQ water were added to achieve to a final volume of 9.5mL. Next, 9.5 mL 2xHBS (50mM HEPES 50 mM, 1.5mM Na2HPO4, 280mM NaCl and in MQ water, pH 7.07) was added and the tube inverted 4 times. After incubation at room temperature for 60 seconds, mixture was added to 165mL of pre-warmed DMEM High Glucose, supplemented with FBS and Pen/Strep (concentrations as described above).

Transfection mixture was left on cells for 12-14 hours and viral production medium (Ultraculture, Lonza BioWhittaker 12-725F) supplemented with 1% Pen/Strep, 1% L-glutamine, 1% 100mM Sodium Pyruvate, 1% 7.5% sodium bicarbonate and 5 mM sodium butyrate) was added 16 hours post transfection. Virus-containing medium was collected 46 and 65 hours post transfection and was filtered through 0.45 μM Millipore low-protein binding filter units (Millipore SCHVU02RE). Viral supernatant was concentrated by first using low speed centrifugation through 100 kDa MW cutoff Millipore Centricon 70 Plus cartridges to condense volume to < 4mL, followed by ultracentrifugation at 45000 rpm (MLS 50 Rotor) for 1.5 hours, to pellet the lentiviral particles. That pellet was resuspended in 25-30µL viral resuspension buffer (VRB), consisting of 20mM Tris pH 8.0, 250 mM NaCl, 10 mM MgCl2 and 5% sorbitol.

For virus titration in a relevant cell type, neural NE4C cells (ATCC, CRL2925™) were cultured on a 6-well plate in MEM (M5650, Sigma) supplemented with 10% FBS and 1% Pen/Strep. On the day of infection, one well was trypsinized and number of cells recorded. 1µL of concentrated virus was diluted in 2mL of MEM and 5, 50 or 500µL were added to the cells. 1µL of concentrated virus was added to a fourth well, the fifth well was left untreated to serve as negative control. 0.1mg/mL Polybrene (H9268, Sigma) was added to all wells in order to increase transduction efficacy and plates were centrifuged for 30min at 1100xg and 37°C. After centrifugation, virus-containing medium was replaced with fresh MEM complete medium, and cells were incubated for 48 hours. For analysis, wells were washed with PBS and trypsinized. Cells were spun for 5 min at 100xg and resuspended in PBS three times. After the third resuspension, cells were passed through FACS tube filters and left on ice until they were analyzed in a flow cytometer to determine infectious titer.

### Brain dissociation with Papain and Flow cytometry analysis

Papain enzyme (P4762, Sigma) was activated for 30 min at 37°C in 1x HBSS, supplemented with 1.1mM EDTA, 0.067mM beta-ME and 5.5mM L-Cystein. For E13 mouse brains, activation solution was diluted with 1xHBSS with 200U/mL DNaseI and 1mM MgCl, to a final concentration of 3U papain/mL. Brains were incubated for 15-30 minutes in a shaking heating block, at 37°C. Tissue digestion mix was spun down for 5min at 300xg and supernatant was discarded. 1 mL of inhibitory solution (1xHBSS with 200U/mL DNaseI and 1mM MgCl) was added and tissue was triturated by slowly pipetting up and down 8 times. After two more rounds of trituration, the cells were resuspended in ice cold PBS and passed through FACS tube filter caps. Solutions were left on ice until they were analyzed in a Canto LSR II. Forward and side scatter was used for gating to leave out cell debris and doublets. 10000 cells were recorded for each sample and between three and seven embryos from two different litters were analyzed.

### Transfection and shRNA knock down efficacy

NE4C cells (ATCC, CRL2925™) were seeded 16 - 20 hours prior to transfection on a 12-well plate in MEM, supplemented with 10% FBS and 1% Pen/Strep. Lipofectamine 2000 (13778150, Invitrogen) and plasmid DNA were diluted in Opti-MEM (31985070, Gibco) to a final DNA:Lipofectmine ratio of 1:3 and incubated at room temperature for 5 minutes. 1µg DNA was added per well and medium was changed after 8 hours incubation. 24 hours post transfection, wells were washed with ice-cold PBS and either fixed in 4% formalin or used for qPCR and Western Blot. RNA was extracted using Qiagen RNeasy MiniPrep kit (74106, Qiagen) and cDNA was produced using RT-PCR kit (K1612, Thermo Scientific). For qPCR, cDNA was mixed with Fast SYBR Green (4385612, Applied Biosystms) and run for 40 cycles. Primer sequences were β-actin F: gacaggatgcagaaggagat; β-actin R: ttgctgatccacatctgctg: Sptbn2 F: cttgggctagtgtggaccat; Sptbn2 R: ccatctctccaactggtggt.

### Cell Profiler

Separate images of DAPI and GFP (488) from specific brain regions were exported to jpeg file format and loaded into cell profiler. An in-house pipeline was created, in which first the cell nuclei were identified as primary objects in both images separately. Diameter range was set between 5 and 15 pixels and for calculating the threshold, Otsu and a two-class thresholding method was used. Finally, the percentage overlap of both images was calculated to obtain the number of GFP+ nuclei per section.

### Cloning

pLKO1 vector containing shRNA (Sigma Aldrich) and H2b-GFP plasmids (gift from Elaine Fuchs; Addgene plasmid #25999) were digested with SacII and SphI enzymes for 15min to 1h at 37°C. Digests were run on a 1% Agarose gel at 100V for 1h. For pLKO1 the middle band at 2243bp (containing the shRNA of interest) and for H2b-GFP the band at 5263bp (containing the H2b-GFP) was excised and DNA was extracted using Qiagen QuickGel extraction kit (28704, Qiagen). Ligations were set up with a molar ratio 3:1 of insert:vector. Using NEB Quick ligation kit (M2200S, Bionordika), ligation mixture was incubated at room temperature for 5 min. The reaction was put on ice and competent cells (Stabl3, C737303, Invotrogen) were transformed following the manufacturers protocol. 100µL transformation mix was spread on an Agar plate containing 100µg/mL Ampicillin (A5354, Sigma) and incubated over night at 37°C. The next day, 3-5 colonies were picked and incubated in 2mL Terrific Broth (T0918, Sigma) containing 100µg/mL Ampicillin for 8 hours. Plasmid DNA was extracted using Qiagen MiniPrep kit (27104). A diagnostic digest was performed using NdeI and SacII in order to discriminate between empty H2b-GFP (290bp + 7261bp) and clones with the shRNA (323bp + 7261bp). Digests were run on 2% Agarose gel at 60V for 3h. Insert-positive vectors were validated by sequencing using the Eurofins barcode sequencing service.

### MiniPromoters

The LV-MiniP-H2B-GFP vectors were prepared by inserting the respective MimiP sequences (*24*) into the LV-H2B-GFP vector (*3*) (Addgene, #25999) using PCR introduced SalI, FseI restriction sites. In detail, hPGK promoter was removed from the LV-H2B-GFP vector by PasI digest, dephosphorylated by CIP (both NEB), and gel purified (QIAGEN, 28706). The promoter-less linear backbone plasmid, and MiniP containing pEMS1172 (Addgene, #29301), pEMS1375 (Addgene, #29176), and pEMS1199 (Addgene, #29100) plasmids (Portales-Casamar et al., 2010), were used as templates for SalI, FseI restriction sites introduction by Phusion Green High-Fidelity DNA Polymerase PCR, using manufacturers protocol (30cycles, 2-step protocol, no GC additive, 5% DMSO) (Thermo Fisher Scientific, F534S), and JM120F/R, JM121F/122R primers. Appropriate PCR products were cut from gel and purified (QIAGEN, 28706). The purified PCR fragments were digested by SalI/FseI restriction enzymes (NEB) and LV-H2B-GFP backbone was also dephosphorylated (CIP, NEB). Resulting DNA fragments were directly column purified (QIAGEN, 28706), ligated with T4 DNA ligase (NEB), and transformed into One Shot™ Stbl3™ Chemically Competent E. coli. Resulting clones were screened by PCR (DreamTaq Green PCR Master Mix, thermo Fisher Scientific; K1081), restriction digest, and Sanger sequencing. Primer sequences were:

JM117F ggtctccttaactgcaccct

JM117R atatgggtcactgaagcgct

JM118F ggcgcacaactgtaattcca

JM118R cctcaacctcccagccttaa

JM119F acagtagcattgcaaagcct

JM119R ggaataagggcaggaaaatatgg

JM120F tatgtactgaGTCGACgaggccctttcgtcttcaa

JM120R tcagatttacGTCGACatggtggcggaagttcctat

JM121F tatgtactgaGTCGACgGGATCTACCATGCCAGAGC

JM122R tcagatttacGTCGACgGCCAAAGTGGATCTCTGCT

## Results

### Establishing conditions for NEPTUNE

Prior to neurulation in mice, around E7.5, the neural plate is exposed to the amniotic fluid and is therefore accessible to injected agents (Fig 1A). Neurulation stages can be clearly discerned by ultrasound until their completion at E9.5 (Fig 1B). Preliminary experiments determined that injections at E8.5 were too late to achieve consistently high levels of transduction since neurulation is ongoing (data not shown). We therefore focused on E7.5 for *in utero* transduction. In order to determine the optimal volume for embryo survival, we injected a range of volumes of viral resuspension buffer (VRB) into the amniotic cavity (ca 20-500nl) at E7.5 and recorded the percentage of injected embryos per litter that survived injections 5 days later at E13 (Fig 1C, D). Sham injections identified a baseline resorption rate of 0 to 25%. Therefore, we drew the cut-off for acceptable survival rates at 70% and identified the highest permissible volume for this survival rate (Fig 1E). However, at E7.5, embryonic development and the size of embryonic cavities can vary greatly, even within litters (Supp Fig 1A). Therefore, the predictor of survival was not volume per se, but rather the relative increase in amniotic volume (Fig 1 F, and data not shown). Ultrasound-checks prior to surgery can be used for embryo staging (Supp Figure 1B).

**Figure 1.**
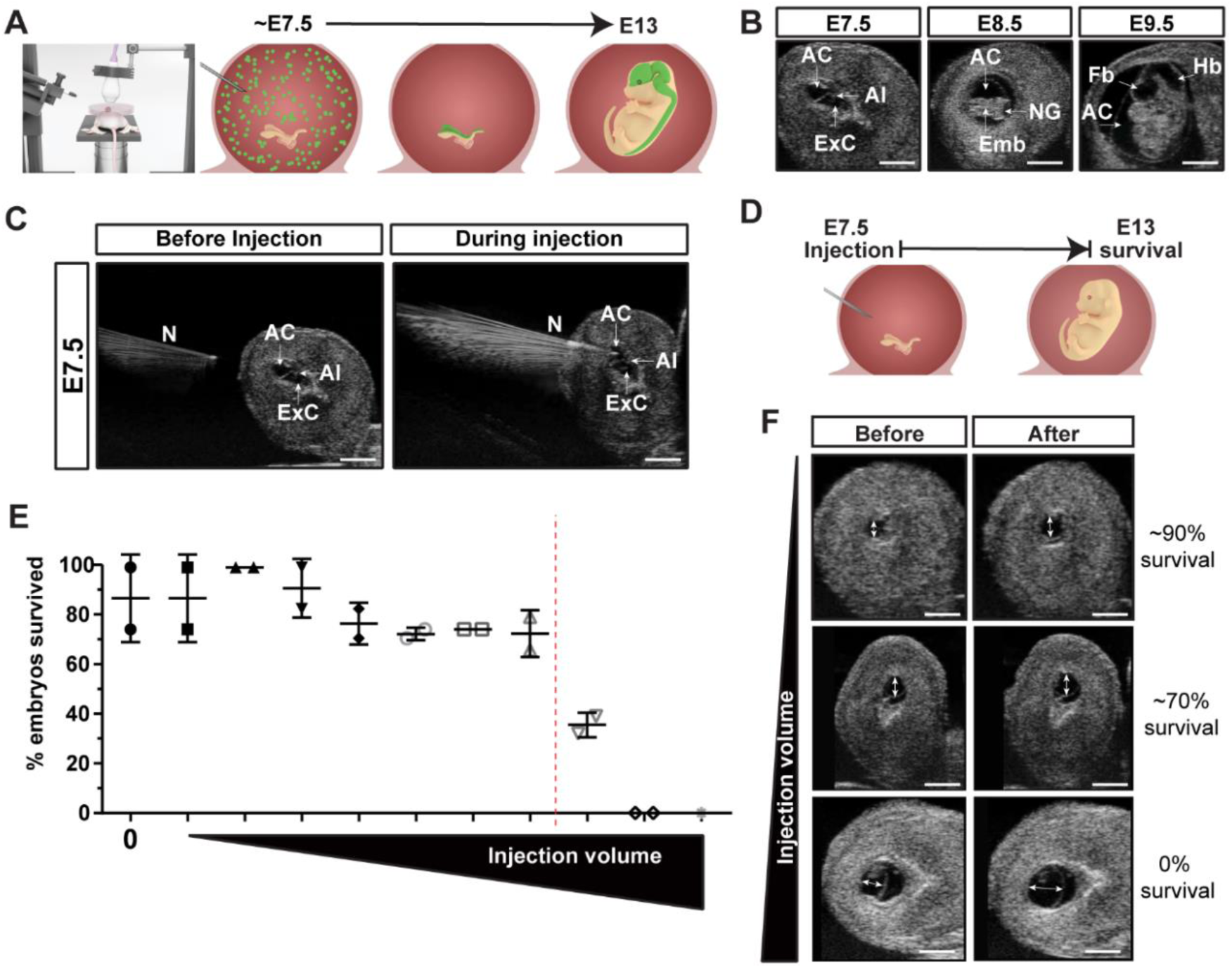
Ultrasound-guided in utero nano-injection is sensitive to volumes injected. (A) At E7.5 the pregnant female is anesthetized and the uterus is externalized. Injection prior to neurulation at E7.5 is expected to target the future nervous system. (B) Neural development between E7.5-E9.5 is easily discerned by ultrasound, allowing identification of the amniotic cavity (AC), allantois (Al), and extracoelomic cavity (ExC) at E7.5 embryo; the embryo (Emb), or neural groove (NG) at E8.5 when neurulation is near completion, and of the neurulated brain at E9.5 with an obvious forebrain (Fb) and hindbrain (Hb). (C) A sharp needle (N) and steady injection ensures the amniotic cavity is injected without puncturing the embryo or allantois. (D) Survival was assessed by injecting viral resuspension buffer (VRB) at E7.5 and collecting embryos at E13.5. (E) Sham injection with no VRB led to 87% survival of embryos. Increasing volumes between 20-500nl of VB were injected, revealing a sharp cutoff in survival at higher volumes which (F) visibly distended the amniotic cavity. Scale bars are 1mm.

### Widespread and stable transduction of the murine CNS upon lentiviral injection at E7.5

Following volume optimization for E7.5 injections, we next determined which viral titers are required to successfully transduce the entire neural plate, using a lentiviral construct that induces stable expression of a H2B-GFP fusion protein (*3*). We used the maximum volume compatible with >70% survival, testing titers by injecting at E7.5, and collecting at E13.5. To ensure an unbiased quantification of transduction efficiency we employed two different techniques to quantify GFP. Half of each brain was dissociated and analyzed using flow cytometry, while the other half was sectioned and stained for GFP and DAPI, and GFP+ nuclei were quantified using Cell Profiler (*25*) (Figure 2A). Tissue collection at different developmental and adult stages confirmed that lentiviral transduction was strong in brain and stable until at least 6 months (Figure 2B, C). Both flow cytometry and Cell Profiler quantification resulted in equally sensitive detection of GFP positive cells. (Figure 2D, E). Using the optimal volume and titer >95% of the cells in the developing CNS could reproducibly be targeted with NEPTUNE (Figure 2D, F).

**Figure 2.**
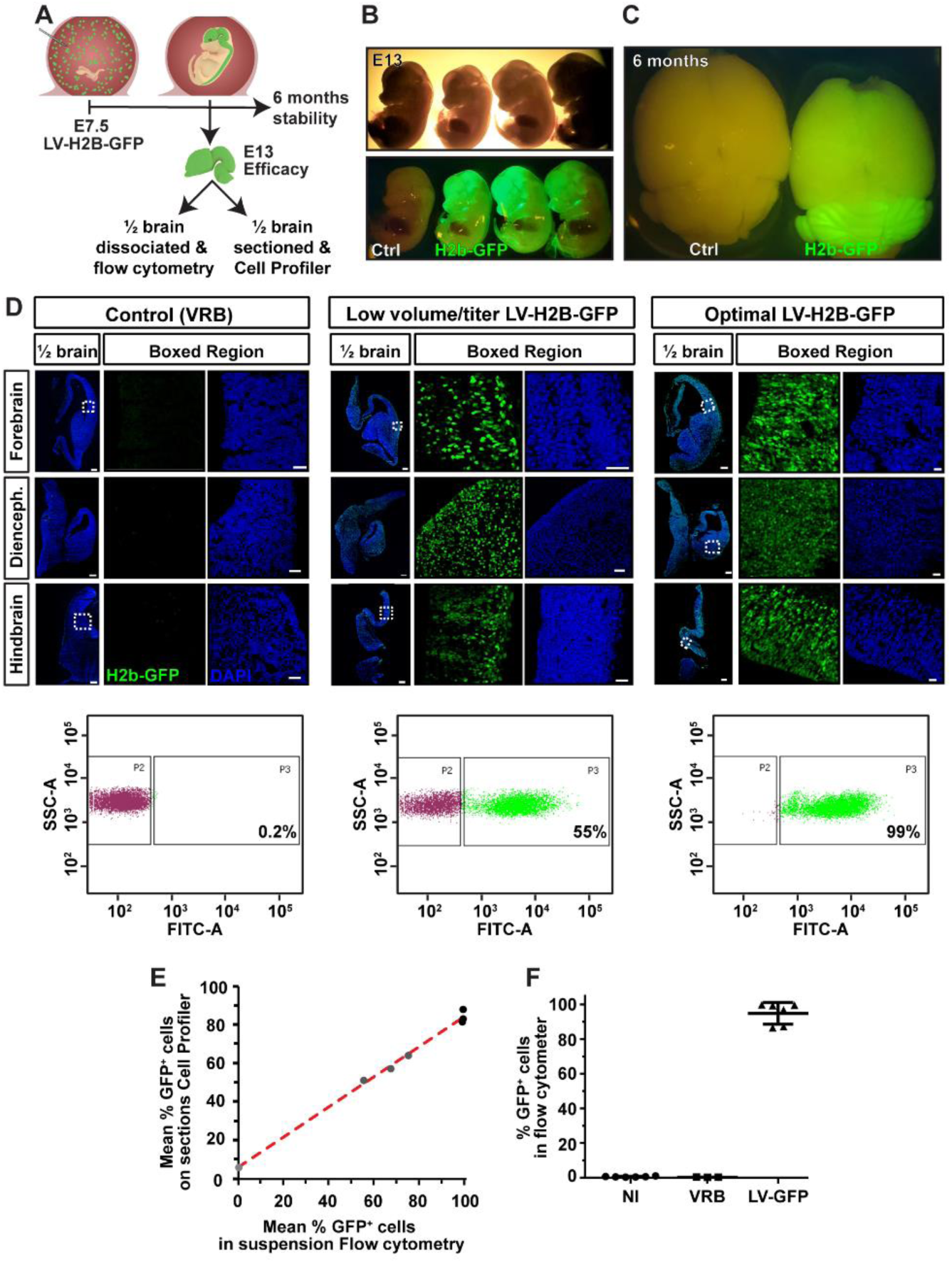
Lentiviral injection at E7.5 results in widespread and stable transduction of the murine CNS. (A) The amniotic cavity was injected at E7.5 and embryos were either allowed to be born and assessed at 6 months, or collected at E13.5. E13.5 brains were divided and half and assessed for GFP positivity using flow cytometry of dissociated cells or CellProfiler analysis of sectioned brain. (B, C) Brain targeting is obvious at E13.5, and stable until 6 months. (D) Low volume/titer injection results in 55% transduction of cells in brain, while optimal volume/titer results in up to 99% GFP+ cells, assessed by flow cytometry. (E) Flow cytometry and CellProfiler quantification of the same brains are well correlated across efficiencies. (F) Optimal volume/titer yields high-efficiency transduction of 85-99%. Scale bars are 200µm (1/2 brain tiles) and 50µm (boxed regions).

### Lentiviral transduction is even across CNS regions and cell types

The neural tube closes at specific neural tube closure points, potentially making these regions less amenable to lentiviral manipulation once these sites have closed. In mice, the initial neural tube (NT) closure point is at the hindbrain/cervical boundary, closure point 2 is at the forebrain/midbrain boundary, and closure point 3 is at the most rostral end of the forebrain. Closure/zippering proceeds rostrally and caudally from closure points 1 and 2, while closure proceeds backwards from closure point 3 towards closure point 2 (*26*). Injections at E8.5 resulted in variable efficiency of transduction, from anterior to posterior brain and spinal cord, reflecting partial neural tube closure (data not shown).

We therefore assessed transduction efficiency throughout the brain and spinal cord, from anterior to posterior, and in neural stem cells, glia or neurons to determine whether the entire nervous system was evenly targeted by E7.5 injections. (Figures 3 and 4). H2B-GFP was highly and evenly expressed from anterior to posterior brain, in both SOX2^+^ neural stem cells and in post-mitotic NeuN+ neurons (Figure 3A,B). H2B-GFP was also widely expressed in the cerebellum (Fig 3C), which develops from a cerebellar primordium first evident at E12.5. H2B-GFP was expressed in both GFP+ Calbindin positive (CALB1+) Purkinje neurons and in SOX2+ glia (Fig 3C).

**Figure 3.**
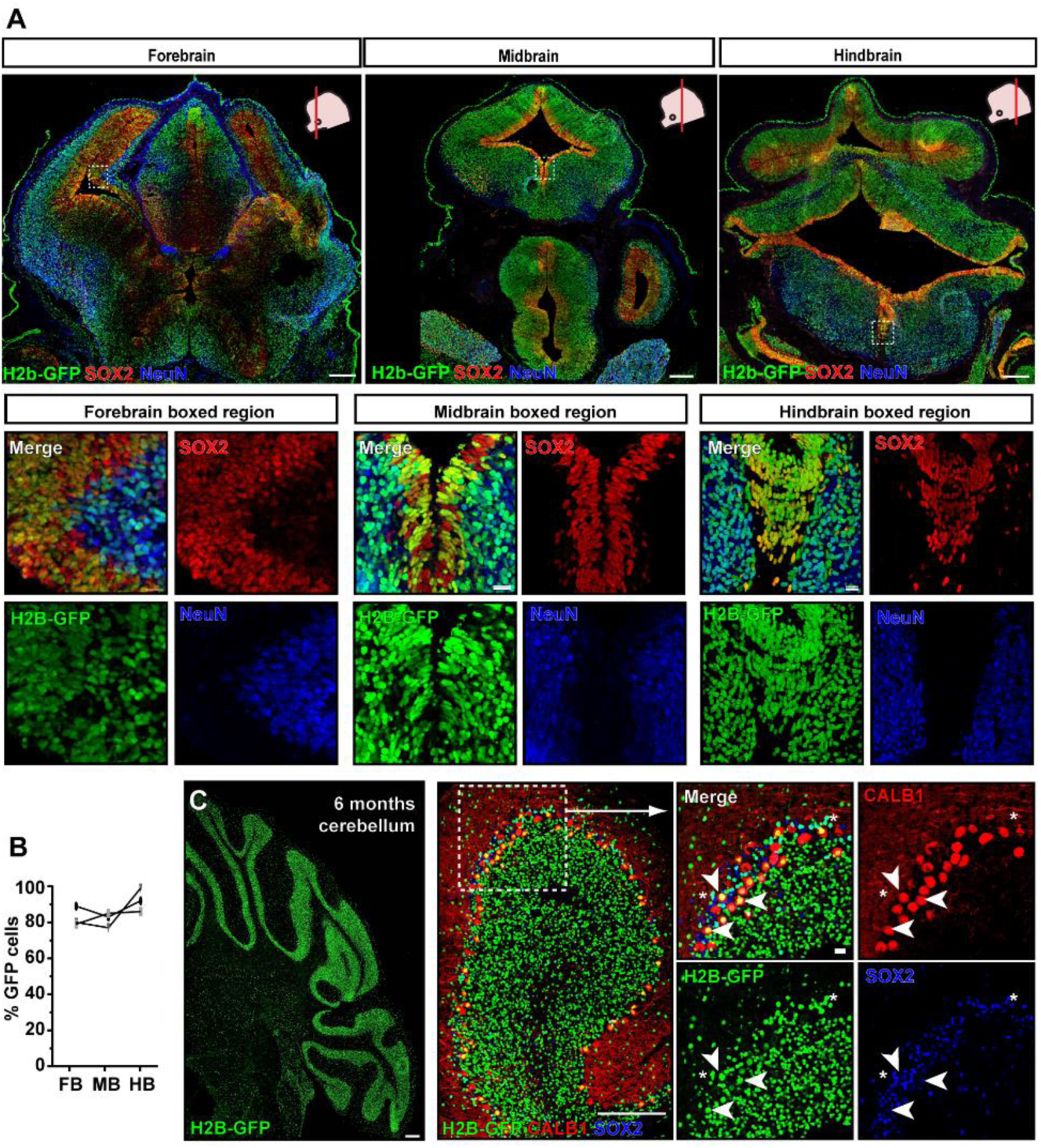
Lentiviral transduction is even across brain and neural cell types. (A) GFP transduction efficiency from anterior to posterior brain, stained for GFP, neural stem cells (SOX2) and neurons (NeuN) shows even transduction across brain and in both cell types. (B) Quantification of anterior-posterior GFP+ cells using CellProfiler confirms even transduction. Scale bars are 200µm (top panels) and 20µm in boxed regions. (C) Transduction at E7.5 also contributes to cerebellum, a brain region that arises later in development, labelling both neurons (CALB1) and glia (SOX2) at 6 months. Scale bars are 200µm (tiles) and 20µm boxed regions.

**Figure 4.**
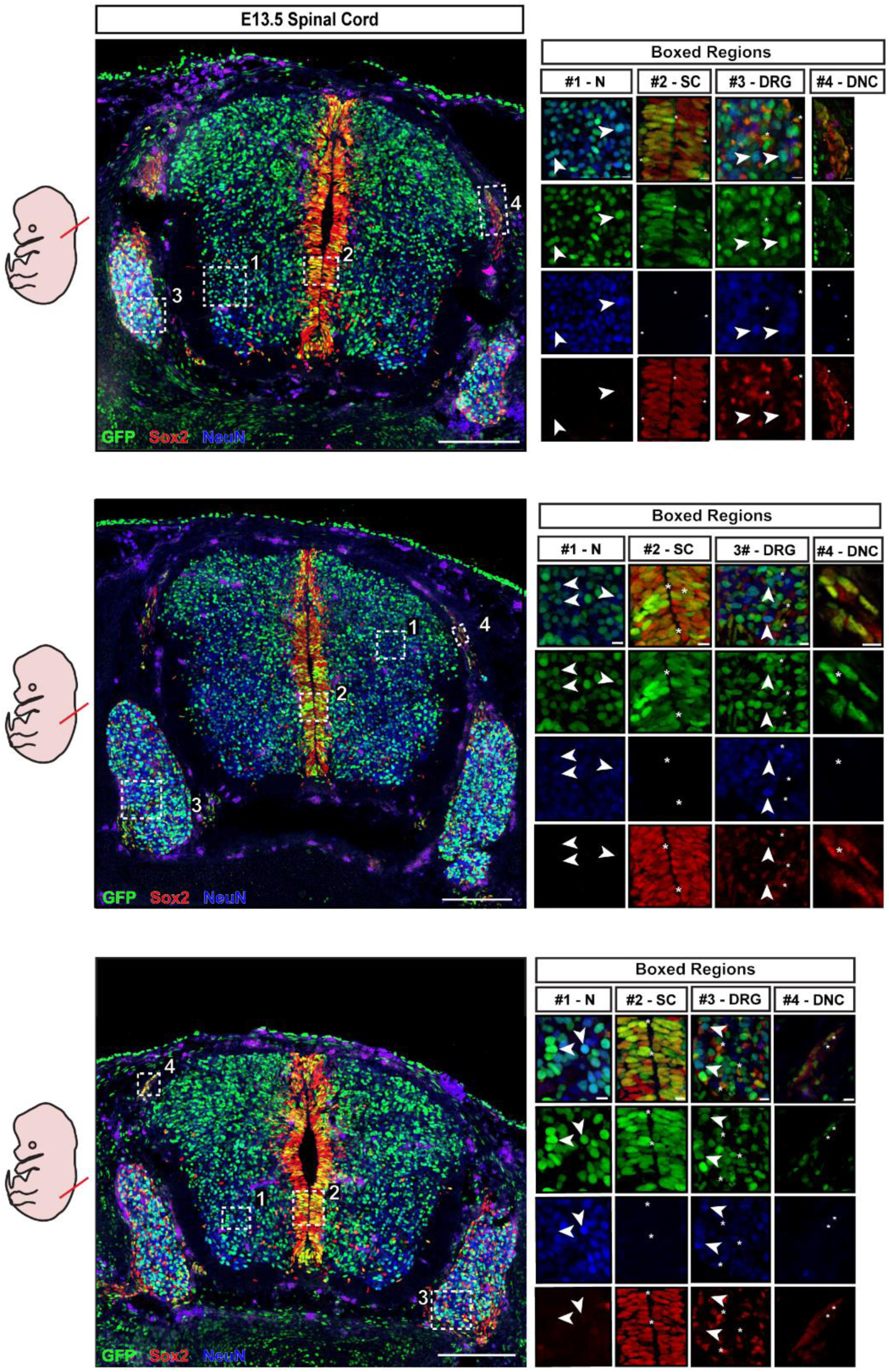
Lentiviral transduction is even across spinal cord and neural cell types. (A) GFP transduction efficiency from anterior to posterior spinal cord, stained for GFP, neural stem cells (SOX2) and neurons (NeuN) shows even transduction across brain and in both cell types. Neuronal populations (N), and neural stem cells (SC) are targeted. Neural crest is also targeted, as evidenced by GFP expression in delaminating neural crest (DNC) and dorsal root ganglia (DRG). Scale bars are 200µm (left panels) and 10µm (boxed regions).

A similarly even transduction pattern was present in the spinal cord (Figure 4). Virus injection at E7.5 efficiently transduced the neural plate contributing to spinal cord from anterior to posterior, and contributed to SOX2+ progenitor cells and NeuN+ mature neurons, as well as delaminating neural crest cells and dorsal root ganglia (Fig 4). Since the virus is delivered into the amniotic fluid and is expected to transduce all cells it comes into contact with, we expected that other organs and tissues would be targeted with this approach. Therefore, we assessed whole body sections at E13.5 and identified highly positive signal in skin, lung and the intestinal tract (stomach, intestine) as well as in the liver and heart (Supp Fig 3A and data not shown), predominantly in cells either in contact with amniotic fluid, such as the lungs or gastrointestinal tract, or which are expected to be derived from neural crest.

### Cell-type specific expression with NEPTUNE and miniPromoters

NEPTUNE targets the entire central nervous system evenly, as well as the neural crest. In order to achieve a more versatile and specific method to investigate gene function in brain, we next aimed to develop conditional gene expression, without the use of dedicated Cre-driver mouse strains. We cloned miniPromotor sequences identified by the Pleiades Promoter Project (*24*) as driving expression in neuronal progenitors (*Doublecortin, Dcx* miniP), astrocytes (*Glial fibrillary acidic protein, Gfap* miniP) and oligodendrocytes (*Oligodendrocyte Transcription Factor 1, Olig1* miniP), and replaced the PGK promoter driving H2B-GFP in *LV-H2b-GFP*, creating instead *Dcx-H2b-GFP, GFAP-H2b-GFP* and *Olig1*-*H2b-GFP* (Figure 5A). *In utero* transduction with the parental *LV-H2b-GFP* at E7.5 led to widespread expression in skin (Figure 5A) as well as in the nervous system (Figure 5B). However, transduction with *Dcx-H2b-GFP* resulted in specific GFP signal from the cortex, and no signal in skin (Figure 5A).

**Figure 5.**
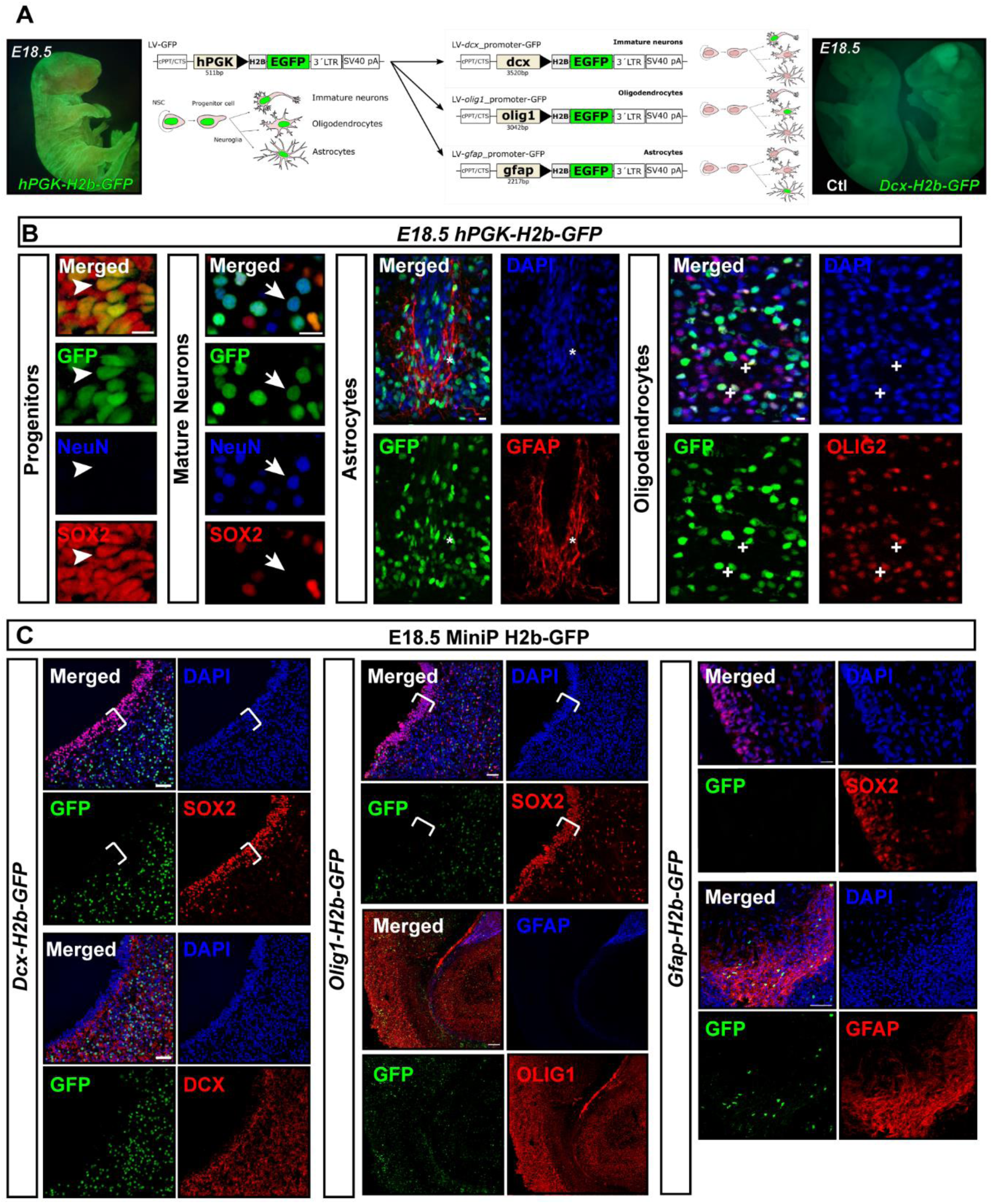
Cell-type specific expression with NEPTUNE and miniPromoters. (A) Transduction with hPGK-H2b-GFP (LV-GFP) at E7.5 results in widespread expression in skin as well as brain. MiniPromotor sequences for Dcx, Olig and Gfap were cloned into LV-H2b-GFP, replacing the PGK promotor, to drive expression in neuronal progenitors, oligodendrocytes or astrocytes. Transduction with Dcx-H2b-GFP resulted in obvious GFP expression in brain, with no GFP+ cells in skin. (B) Transduction with hPGK-H2b-GFP at E7.5 targets future progenitors (SOX2+), neurons (NeuN+), astrocytes (GFAP+) and oligodendrocytes (OLIG2+) at E18.5. Scale bars are 10µm. (C) MiniPromotors achieve conditional expression. None of the constructs was expressed in SOX2+ neural stem cells (white brackets). Dcx-H2b-GFP is exclusively expressed in neuronal cells, Olig1-H2b-GFP is widespread in OLIG1 positive regions, and Gfap-H2b-GFP was exclusively expressed in GFAP+ cells. Scale bars are 50µm.

We confirmed that viral transduction at E7.5 with *LV-H2b-GFP* contributes to SOX2+ neural progenitors, NeuN+ neurons, GFAP+ astrocytes and OLIG2+ oligodendrocytes at E18.5 (Fig 5B). In contrast, *Dcx-H2b-GFP* was not expressed in SOX2+ neural progenitors at E18.5 (Fig 5C, left-hand panels, white brackets), but was expressed in DCX+ neurons (Figure 5C, lower panels). *Olig1*-*H2b-GFP* was also not expressed in the SOX2+ neural progenitors (Fig 5C, middle panels, white brackets), but was widespread in OLIG1+ regions. Finally, *GFAP-H2b-GFP* was also negative in SOX2+ neural progenitors (Fig 5C, right-hand panels), and was instead expressed in GFAP+ astrocytes (Figure 5C, right-hand lower panels). In conclusion, NEPTUNE achieves >95% transduction of all cell types in brain with the PGK promotor, and expression can be directed to specific cell types using miniPromotors.

### NEPTUNE reveals a novel role for Spbtn2 in neurulation

Next, to test whether NEPTUNE is able to induce nervous-system wide gene dysregulation with consequences at the whole-organ level, we designed proof of principle experiments focusing on a neurodevelopmental defect. The *Spectrin Beta, Non-Erythrocytic 2* (*Sptbn2*) gene is highly expressed in the CNS and holds many essential functions in intracellular vesicle transport and cytoskeleton dynamics (*27*). Furthermore, mutations in *SPTBN2* are associated with both Spinocerebellar ataxia 5 (SCA5, OMIM 600224) (*12*–*16*) and spinocerebellar ataxia, autosomal recessive 14 (SCAR14, OMIM#615386) (*17*–*19*). Mouse models for *Sptbn2* mutation display an adult-onset defect in motor function, but these mice express shorter SPTBN2 isoforms (*21*) or alternative splice-variants (*22*), suggesting that this phenotype may not reflect complete loss of function. We therefore decided to test whether knock-down of *Sptbn2* at E7.5 induced an ataxic phenotype or a more severe phenotype, as would be expected if the splice-variants or shorter variants were compensating for loss of full-length SPTBN2.

Five shRNAs against *Sptbn2* were tested in the neural cell line NE4C, revealing similar 50-60% downregulation of *Sptbn2* mRNA levels for all constructs, compared to scrambled control, at 24 and 48 hours (Figure 6A). After cloning two of the shRNAs into LV-H2B-GFP (#1 and #2), knockdown efficacy was maintained (Figure 6B). The 50% knockdown is likely due, at least in part, to the difficulty of transfecting NE4C cells, which showed circa 50% transfection rates (Figure 6 C).

**Figure 6.**
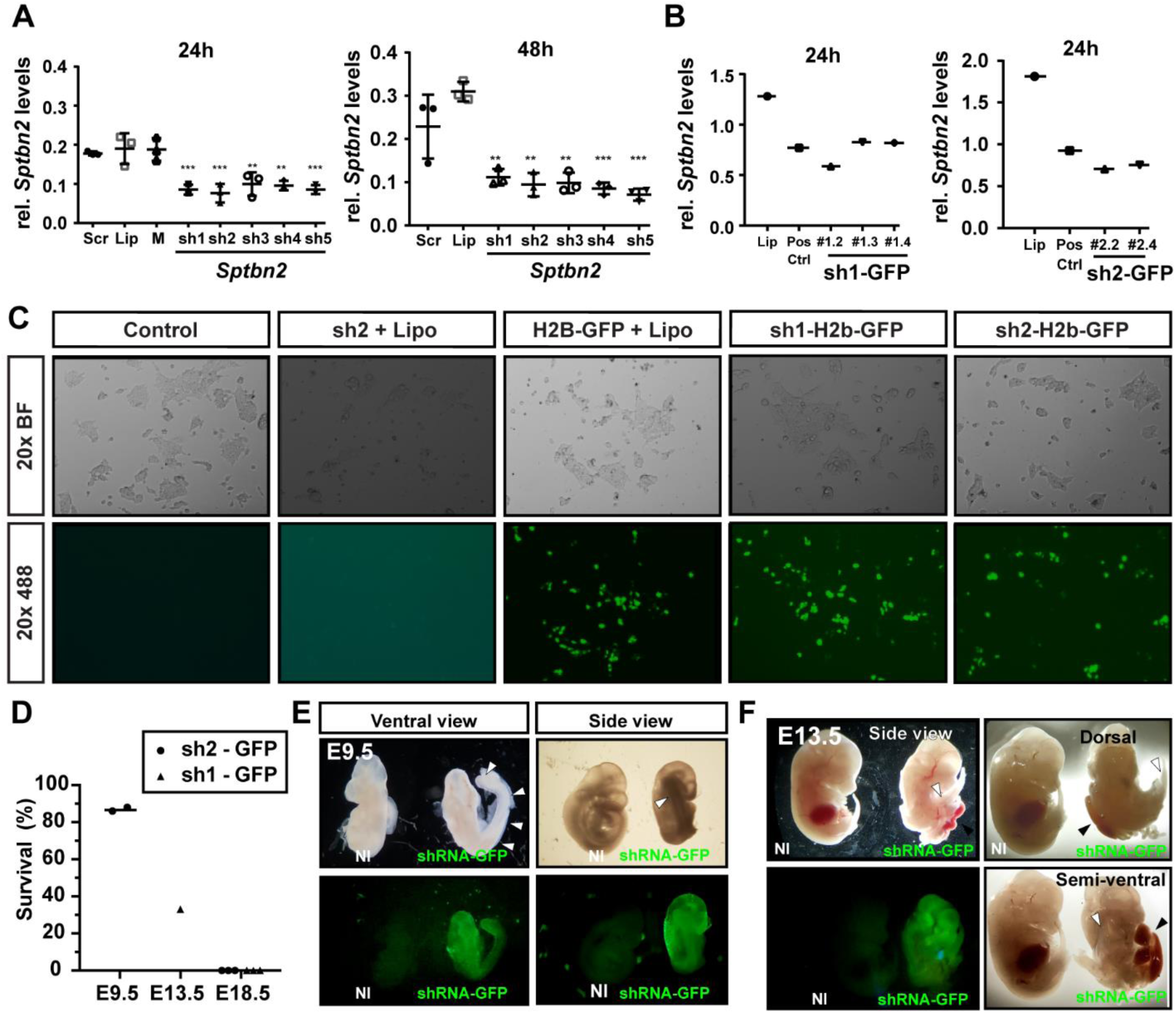
Knockdown of Sptbn2 in vivo with NEPTUNE reveals a novel role in neurulation and abdominal wall closure. (A) Five shRNAs against Sptbn2 were tested for knockdown efficiency in neural NE4C cells at 24 and 48 hours. (B) shRNA #1 and #2 were selected for subcloning into hPGK-H2b-GFP, and maintained knock-down efficiency was confirmed for several clones after 24 hours transfection in NE4C cells. (C) Bright field and fluorescence imaging of transfected NE4C cells confirmed GFP expression for sub-cloned shRNAs and revealed ca 50% transfection efficiency. (D) In utero transduction of embryos with shRNA#1 or shRNA#2 against Sptbn2 resulted in 87% survival at E9.5, 33% survival at E13.5 and 0% survival at E18.5. (E) At E9.5 injected embryos displayed a neurulation/turning defect with rightward skewing of the body axis (white arrowheads). (F) At E13.5, embryos continued to display halted turning, rightward skewing of the body axis (white arrowheads) and abdominal wall defects with externalized abdominal organs (black arrowheads).

We injected 9 litters of mice with virus encoding *Sptbn2* shRNA (construct #1 or #2) at E7.5 and collected embryos at E9.5, E13.5 and E18.5. Surprisingly, no embryos were recovered at E18.5 (6 litters, 3 injected with shRNA#1, and 3 injected with shRNA #2, total 41 embryos, Figure 6D). At E9.5, 87% of injected embryos survived (2 litters, 15 injected embryos injected with shRNA #2, Figure 6D), but displayed strong skewing of the body axis and tail to the right (Figure 6E, white arrowheads) with a shortened right-hand side of the body, longer left-hand side, arrested turning, and were smaller (Figure 6E). At E13.5 only 33% of injected embryos survived (1 litter, 6 embryos injected with shRNA #1, Figure 6D), and the phenotype had progressed with strong rightward skewing (white arrowheads), failed turning and an open abdominal wall with an externalized liver (black arrowheads) (Figure 6F). Survival rates dropped greatly between E9.5 to E13.5, with 0% survival at E18.5, suggesting that the absence of *Sptbn2* is embryonic lethal and crucial for neural tube development and abdominal wall closure.

## Discussion

Although electroporation of chick and mouse are powerful tools for studying gene function in the nervous system, traditional mouse genetics have been the gold standard for testing gene function *in vivo*, due to mosaic effects in electroporation studies.

Ultrasound-guided *in utero* nanoinjection has proven to be a powerful tool to unravel complex genetic networks in skin (*3*–*5*). Now, we show that the developing mouse nervous system can be targeted in a highly efficient and reproducible manner, achieving over 95% transduction efficiency throughout the brain and spinal cord (Figures 3 & 4). Transduction is stable to adulthood (Figures 2 and 3). Introducing the miniPromoter sequences enabled expression in defined cell types, without the need for transgenic *Cre* mouse lines (Figure 5). Finally, we demonstrated that knock down of *Sptbn2 in vivo* results in severe neural tube defects, revealing a completely new function for this gene in neurulation (Figure 6).

NEPTUNE allows high efficiency and wide-spread transduction of the nervous system. We identified volumes injected, viral titer, and freshness of the virus as crucial determinants of transduction and survival efficiency (Figure 1, 2, and data not shown), and optimized these to achieve >95% transduction in brain and spinal cord. In order to achieve conditional expression, we replaced the PGK promotor with miniPromotors (*24*) for *Dcx, GFAP* and *Olig1*. These promotors showed no leakiness in SOX2+ precursor cells (Figure 5C), and labelled specific cell types (neurons, astrocytes and oligodendrocytes), albeit with somewhat variable efficacy. While the *Dcx-miniP-H2B-GFP* construct was active in neurons (Fig 5C), the *Olig1-miniP-H2B-GFP* could be found in OLIG1 negative cells. This may reflect transient *Olig* expression in glial precursors contributing to astrocytes and oligodendrocytes, in which expression is downregulated during differentiation into astrocytes, while the EGFP reporter is stabilized via fusion to H2B and would be expressed longer. In order to circumvent induction of a reporter by transient expression, future modifications could include introduction of a floxed-STOP cassette before H2B-GFP, and using constitutive Cre-ERT2 mice, allowing tamoxifen-induced excision of the STOP cassette at the desired time-point and expression only in cells currently expressing the promotor of interest.

Further future modifications and improvements to NEPTUNE could additionally increase its versatility. Transduction of virus encoding sgRNA into Cas9-expressing mice would allow targeted knockouts or gene editing, either to introduce specific disease-associated mutations, or to correct mutant alleles and test therapeutic gene editing. The rise of AAV evolution (*28, 29*), and thus engineering viral tropism, may also allow targeted transduction of specific organs during embryonic development.

The development of NEPTUNE can reduce the number of mice used in research dramatically. It has been estimated that of 3,872 targeted genes in the mouse, 45% are not essential for viability or fertility (*30*). Conversely, 14% of mutated genes lead to embryonic lethality, usually at or before mid-gestation (*30, 31*). Combining these two, the probability of knocking out a gene, and obtaining knockout mice to study that exhibit an interesting and important phenotype during embryogenesis, is around 40%. The development of NEPTUNE would allow screening for relevant neural phenotypes and circumvents the risk of severe embryonic lethality in heterozygous mice. Furthermore, NEPTUNE could be used to investigate genetic redundancy and dissect apart signaling networks. In this regard, this would also save large amounts of mice. As an example, breeding triple heterozygous mice to obtain triple knockouts would generate 64 pups to obtain one wild type and one triple knockout, creating an excess of 62 unused pups. Assuming one would also study the single and double knockouts as controls, one could use 8 of the 64 pups, nevertheless generating 56 excess pups, meaning 87.5% of mice are not used. With NEPTUNE, it would thus be possible to directly generate the genotypes/knockdowns of interest and not create unused mice. Furthermore, validation of gene function across different strains of mice is facilitated since back-crossing would not be required – further increasing the possibility to verify reproducibility.

The *Sptbn2* phenotype (Figure 6) is more severe than expected based on our current knowledge of *SPTBN2* in the human ataxic syndromes SCA5 and SCAR14. However, given its role as a cytoskeletal component linking actin and the cell membrane (*27*), it is perhaps less surprising that disruption of *Sptbn2* would lead to dysfunction in neurulation and morphogenesis. Importantly, in most patients, the described mutations are missense or in-frame deletions, suggesting remnant protein may be sufficient to execute some SPTBN2 functions (*12*– *19*). Likewise, the *Sptbn2* mutant mice still express slightly shorter forms of SPTBN2, suggesting some functional rescue (*21, 22*). Our data, using two different shRNAs, suggest that *Sptbn2* also has a crucial role in neurulation and embryonic turning. The phenotype mimics planar cell polarity mutants with single or compound mutations in *Scrib, Celsr1* or *Vangl2* (*32*), suggesting that Spectrin may play a key role in mediating planar cell polarity programs during neurulation and turning.

In sum, NEPTUNE is a powerful new technique to modulate gene expression during embryonic development. It can achieve wide-spread, stable and conditional expression in the brain and spinal cord, and can be used to discover new roles for genes in crucial embryonic processes.

## Acknowledgements

We thank Bettina Semsch for expert care of mice and support; Florian Salomons, Göran Månsson and Shigeaki Kanatani from Biomedicum Imaging Core (BIC) for assistance with image acquisition and consultation; Mattias Karlen for Illustrations; Slobodan Beronja, Jozef Vecera and Linus Christerson for helpful discussions and scientific input. We thank Slobodan Beronja for advice and help, and Ellen Ezratty for sharing space and protocols during ER Andersson’s visit to Rockefeller University. Nicole Stokes, Lisa Polak, Ellen Wong at Rockefeller University for excellent help and training when ER Andersson visited Rockefeller. We thank Mary Beth Hatten for also providing space and resources for ER Andersson’s visit.

We thank the following funders for their support of this project: The Swedish Research Council, Karolinska Institutet (KI Foundations, Career Development Grant, PhD student KID funding, and SFO StratNeuro funding, the Center of Innovative Medicine), The Ollie and Elof Ericssons Foundation, the Tornspiran Foundation, the Jeansssons Foundation, Sven and Ebba-Christina Hagbergs Prize and research Grant, Knut and Alice Wallenberg Project Grant, Fredrik and Ingrid Thurings Foundation, Lars Hiertas Minne, The Childhood Cancer Foundation (Barncancerfonden), The Åhlen Foundation, Åke Wibergs Foundation, Tore Nilssons Foundation, and the Swedish Foundations Starting Grant (ERA). We thank the EASL Sheila Sherlock fellowship (JM). Support for this work also came from National Institutes of Health R01-AR27883 and R0-AR050452 (to E.F.).

## Supplementary Figures

**Supplementary Figure 1.**
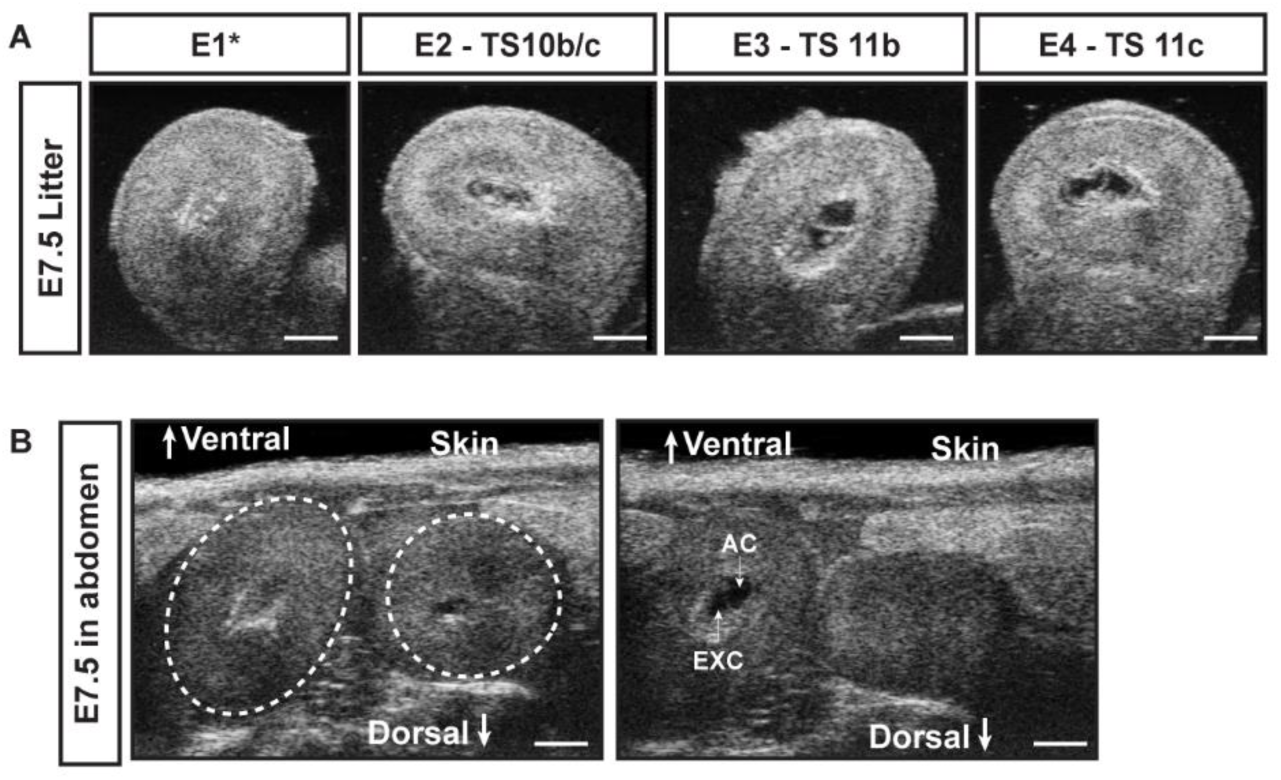
Embryonic staging using ultrasound. (A) Ultrasound images of externalized uterus at E7.5. Gestational age can vary within a litter, and can be observed via ultrasound to determine whether the stage is appropriate for injection. Theiler Stage (TS) 11 was most suitable for high efficiency and high survival. (B) Staging can be assessed prior to surgery, here ultrasound through the abdominal wall of the pregnant female at E7.5 to assess suitability of proceeding with experiment.

**Supplementary Figure 2.**
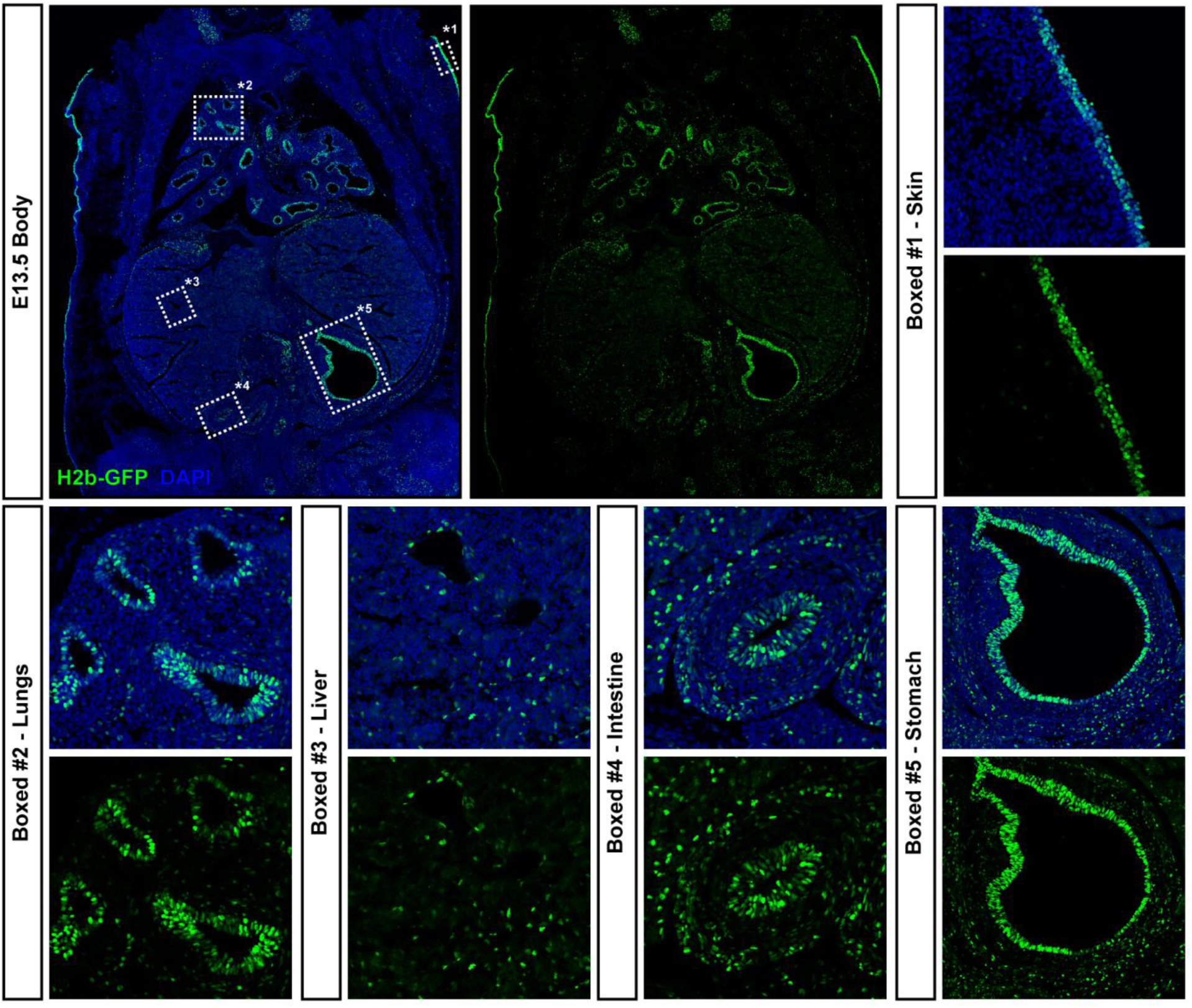
In utero injection at E7.5 targets other tissues in addition to the nervous system. Coronal section of whole abdomen at E13.5shows strong GFP expression is detected in cells in contact with amniotic fluid, or which are expected to derive from neural crest. These include the skin, lungs, stomach and intestine. Liver also contains GFP+ cells, though with a different and diffuse pattern.

